# Limiting factors for queen conch (*Lobatus gigas*) reproduction: A simulation-based evaluation

**DOI:** 10.1101/2021.04.23.441087

**Authors:** Nicholas A. Farmer, Jennifer C. Doerr

**Author notes:** Corresponding author (NF). These authors contributed equally to this work.

## Abstract

Queen conch are among the most economically, socially, and culturally important fishery resources in the Caribbean. Despite a multitude of fisheries management measures enacted across the region, populations are depleted and failing to recover. It is believed that queen conch are highly susceptible to depensatory processes, impacting reproductive success and contributing to the lack of recovery. We developed a model of reproductive dynamics to evaluate how variations in biological factors such as population density, movement speeds, movement restrictions, rest periods between mating events, sexual facilitation, and perception of conspecifics affect reproductive success and overall reproductive output. We compared simulation results to empirical observations of mating and spawning frequencies from conch populations in the central Bahamas and Florida Keys. Our results confirm that low probability of mate finding associated with decreased population density is the primary driver behind observed breeding behavior in the field, although additional factors also play important roles. In particular, sexual facilitation and perception of conspecifics may explain observed lack of mating at low densities and differences between mating frequencies in the central Bahamas and Florida Keys, respectively. Our simulations suggest densities greater than 200 adults/ha are needed for high levels of spawning output, supporting the suggestion that effective management strategies for queen conch should aim to protect high-density reproductive aggregations and critical breeding habitats.

## Introduction

The queen conch (*Lobatus gigas*, formerly *Strombus gigas*) occurs throughout the Caribbean Sea, in the Florida Keys and Gulf of Mexico, and around Bermuda. They are a mollusk characterized by a large, heavy, whorl-shaped shell with multiple short spines at the apex, a brown and horny operculum, and a pink interior of the shell lip [1]. Shell morphology is influenced by environmental conditions including habitat [2, 3]. Females are generally slightly larger than males [4]. They are benthic-grazing herbivores that feed on diatoms, seagrass detritus, and various types of algae and epiphytes [5, 6]. Adult distributions are heavily influenced by food availability and fishing pressure; in unexploited areas, they are most common in shallow marine waters less than 30 m depth [3]. Adults prefer sandy algal flats, but are also found on gravel, coral rubble, smooth hard coral, and beach rock bottoms [7-9].

Adult conch have a protracted spawning season of 4-9 months, with peak spawning during warmer months [1, 10, 11]. They reproduce through internal fertilization, meaning individuals must be in contact to mate. Copulation has been documented day and night [1]. Females can store fertilized eggs for several weeks [10], and egg masses may be fertilized by multiple males [12]. Egg laying takes 24-36 hours, with each egg mass containing about 750,000 eggs [13]. Fecundity appears to be influenced by food availability; with adequate food, females lay an average of 13.6 egg masses during a single reproductive season, compared to an average of 6.7 egg masses containing 500,000 eggs each when food is limited [13].

Queen conch are relatively slow moving, averaging only a few meters of movement per day [14, 15]. Movement rates increase and are fastest in the summer, possibly due to warmer waters promoting increased metabolic activity as well as increased movement related to mate seeking during the reproductive season [14]. In many locations, adult conch migrate to different habitat types during their reproductive season, and then return to feeding grounds [5, 14, 16, 17]. Geographically isolated conch in Florida and Puerto Rico remain in deep water year-round [14, 18].

Queen conch are among the most economically, socially, and culturally important fishery resources in the Caribbean [19, 20], with extremely high domestic and exported landings [21]. The fishery consists of both industrial and artisanal fleets and encompasses the entire Caribbean. Commercial exports increased in the 1980s and 1990s, with a peak of around 3000 tonnes in 1996 and 1997 [21]. These increased landings were accompanied by decreasing population densities across the range [22, 23]. Management approaches vary across the region, but include size restrictions, closed seasons, harvest quotas, and/or gear restrictions. Despite these management interventions, many populations have not recovered [24-26]. In the United States, overharvesting and habitat loss precipitated the collapse of large commercial and recreational fisheries in south Florida. Despite closure of the commercial fishery in 1976, followed by closure of the recreational fishery in 1986, the population has not recovered [27, 28]. NOAA Fisheries initiated an Endangered Species Act status review for queen conch in 2019 to evaluate threats to the species’ habitat, overutilization, and the adequacy of existing regulatory mechanisms to protect the species from extinction [29].

Empirical observations have suggested mating and egg laying in queen conch is directly related to the density of mature adults [30-32]. In animals that aggregate, low population densities can make it difficult or impossible to find a mate [30, 33-35], an issue which is likely compounded for slow-moving animals such as conch [14, 15]. Observations of queen conch populations also suggest an Allee effect, where little to no mating occurs below a critical density threshold [30, 32, 36].

In this study, we present a simple model of conch movement and reproductive behavior to evaluate how well the empirical observations of mating and spawning frequency are explained by variation in density, movement speeds, restricted movement, rest periods between mating events, sexual facilitation, and perception of and/or attraction to conspecifics. This approach provides a stochastic, quantitative approach towards evaluating the relative contributions of a myriad of factors to individual reproductive success and overall population reproductive output. Comparing emergent properties [37] from individual-based mechanistic simulations to empirical observations can benefit hypothesis elimination and facilitate identification of the biological processes driving queen conch reproductive success.

## Materials and Methods

Male and female conch movements were simulated in R [38] using package ‘*particles’* [39]. Sexually mature adult conch were randomly distributed at specified densities on a 1-ha grid and tracked over one-day time steps (Fig 1). Daily reproductive dynamics were simulated across a three-month (92 d) spawning season. The number and frequency of mating and spawning events was tracked and averaged across individuals. Simulation scenarios evaluated the influence of density, movement speed, scent tracking, barriers to movement, interbreeding rest period, sexual facilitation, and conspecific perception distance (Table 1). Nearly all combinations of variables were tested (S1 Data) using custom written R software (S2 File).

**Table 1.** Queen Conch Reproductive Parameters.

| Parameter | Symbol | Units | Value | Source(s) |
| --- | --- | --- | --- | --- |
| Adult Density | N | N ha<sup>-1</sup> | 10, 25, 50, 100, 150, 200, 250, 500, 1000 | n/a |
| Movement Speed | V | M d<sup>-1</sup> | Low: N<sup>a</sup> [2.57 ± 1.57 (1.47 – 3.56)] | Glazer et al. [14], Doerr & Hill [15] |
|  |  |  | Med: N<sup>a</sup> [4.17 ± 0.41<br>(0 – 5.52)]<br>High: N<sup>a</sup> [11.36 ± 0.24<br>(0 – 21.24)] |  |
| Scent Tracking | T | n/a | 0<br>1 (22 m max taxis distance) | Davies & Blackwell [40],<br>Ng et al. [41], Doerr & Hill [15] |
| Barriers to Movement | B | n/a | none, one, long, pylon, cross, horseshoe | n/a |
| Interbreeding Rest Period | RP | d | 0<br>U<sup>b</sup> (0-2)<br>N<sup>a</sup> [8.7 ± 4.9 (0 – 24.83)] | Weil & Laughlin [11] |
| Sexual Facilitation | SF | # prior interactions | 0, 5, 10, 25 | Appeldoorn [33],<br>Gascoigne & Lipcius [54], Delgado & Glazer [36] |
| Perception Distance | PD | m | U<sup>b</sup> (0,0.5)<br>U<sup>b</sup> (0,1.5)<br>U<sup>b</sup> (0,3.0) | Informal expert elicitation from A. Stoner, G. Delgado, R. Glazer |

Table showing the parameters evaluated in simulation models.
<sup>a</sup>Truncated normal distribution [Mean ± SD (Min – Max)]
<sup>b</sup>Uniform distribution

**Fig 1.**
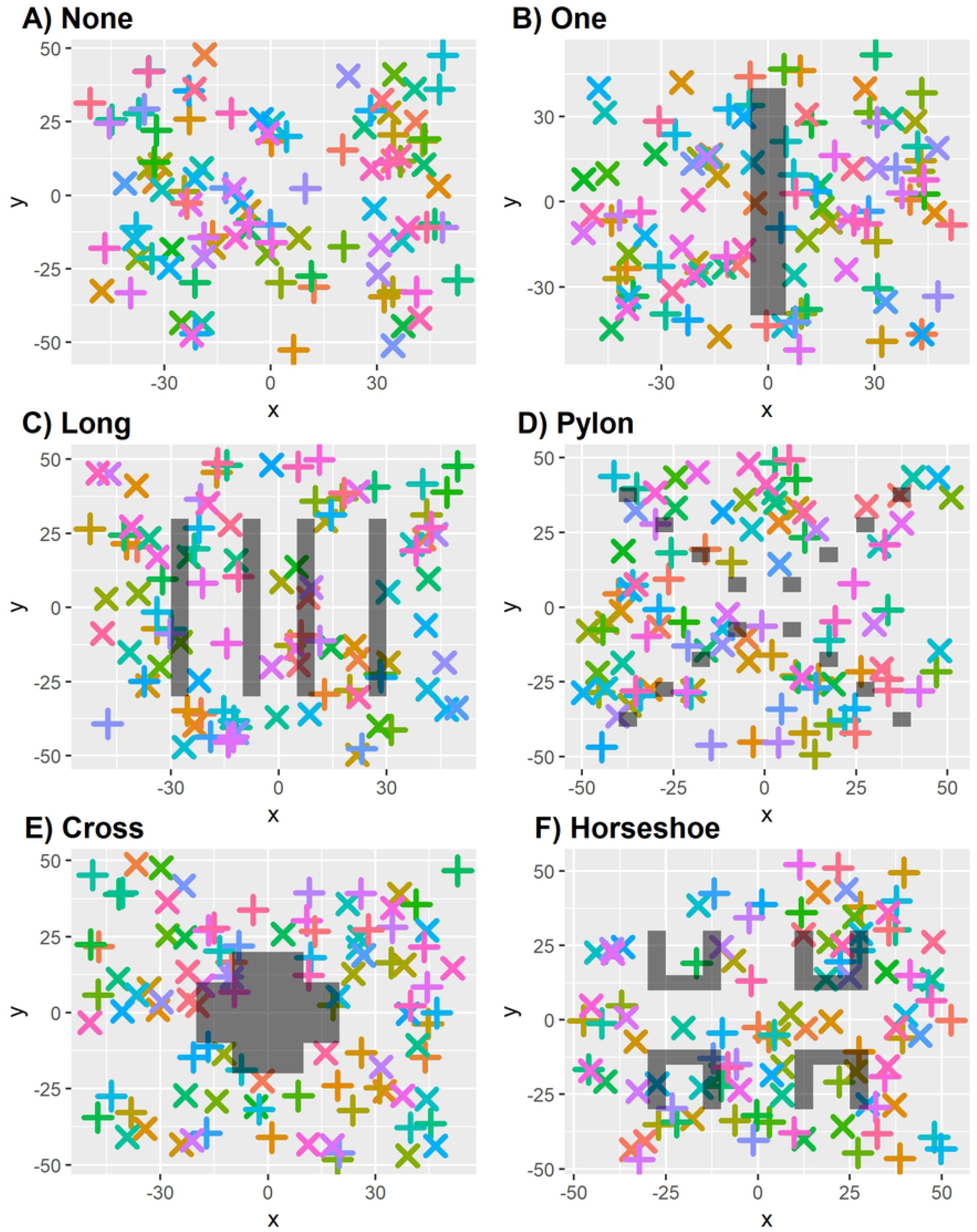
Plot of Barriers to Movement. Barriers to movement (shaded polygons) relative to movements of male (+) and female (x) conch.

Daily movement speeds have been estimated through acoustic telemetry [14, 15]. Adults move at varying speeds throughout the year with movement rates increasing during seasonal migration and slowing during foraging activities or upon reaching mating aggregations. We selected a low speed scenario from male movement speeds and a medium speed scenario from summer movement speeds reported by Glazer et al. [14]. We selected a high movement speed scenario from pre-aggregation migratory movements reported by Doerr & Hill [15]. Movements were simulated as a constrained velocity randomly drawn from a distribution (Table 1) for each day and individual. Movement directions were random unless animals encountered a barrier or scent-trail following was enabled.

Scent-trail following has been postulated as an energy-saving mechanism in gastropods [40, 41]. Because queen conch do not move using a slime trail, it is unclear if scent-trail following is possible for the species. If implicated, it might be accomplished through sex hormone tracking [42]. For most simulations, scent-trail following was disabled. When enabled, scent-trail following was modeled as particle attraction using the *manybody_force* function at a specified level of taxis distance and strength. When enabled, the strength, distance of influence (22 m), and duration of scent-trail following (24 hr) were constrained to reflect the pragmatic constraints of scent trail decay in a dynamic marine environment.

Movement barriers were simulated to evaluate the impacts of microhabitat features on reproductive dynamics (Fig 2). Features ranged from single linear barriers to multiple complex barriers. If a conch’s random movements took them into a barrier, they were returned to the exterior of the barrier and randomly moved from that point.

**Fig 2.**
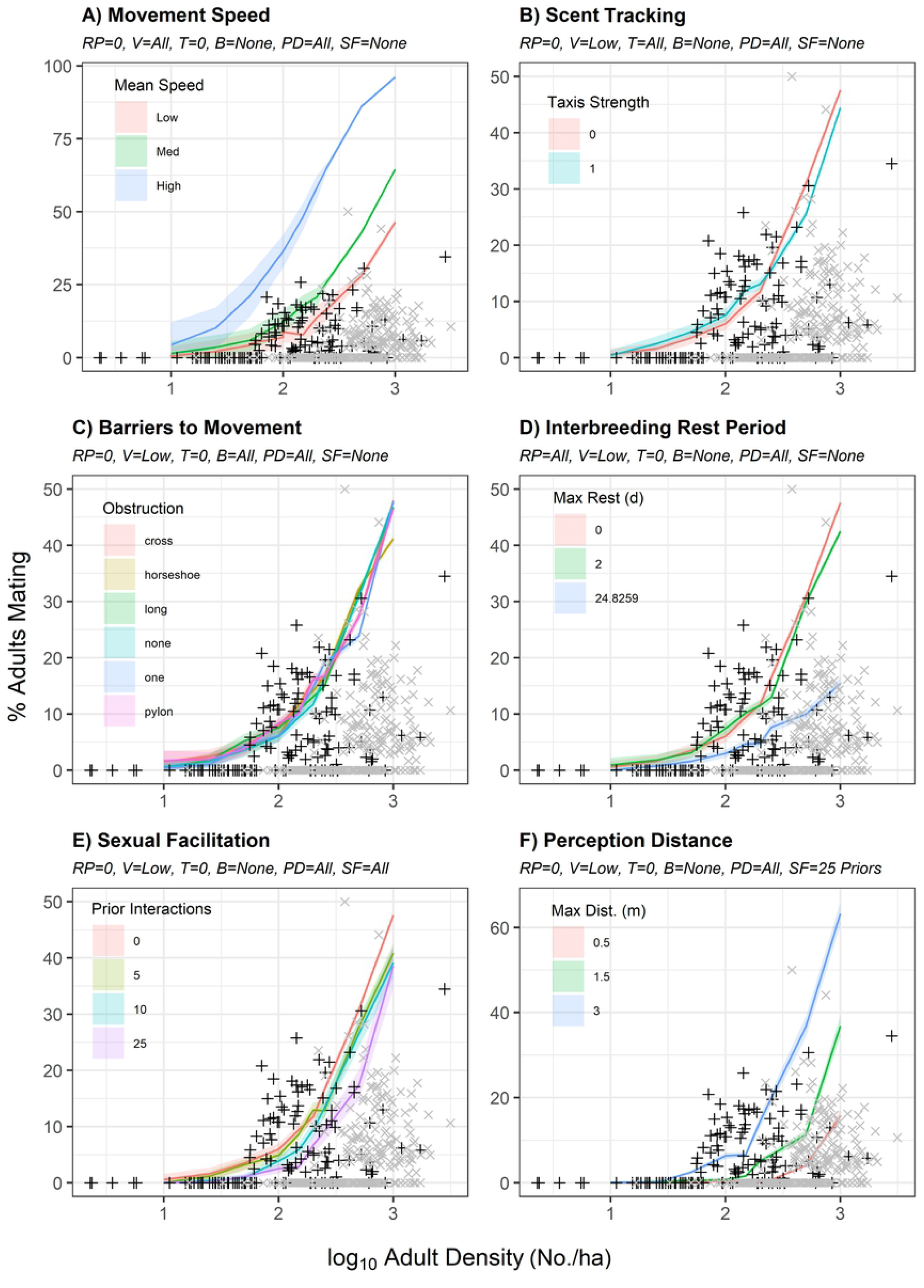
Percent Mating Relative to Model Parameters. Mean (solid line) and 95% confidence band (shaded ribbon) for percent of simulated adult queen conch successfully mating relative to log_10_ adult density (N/ha) relative to variation in A) movement speed (V), B) scent tracking (T), C) barriers to movement (B), D) interbreeding rest period (RP), E) sexual facilitation (SF), and F) perception distance (PD). Empirical observations by Stoner et al. [32] (black crosses) and Delgado & Glazer [36] (gray x’s) are overlaid for comparison.

Female receptivity to mating was evaluated as a possible interbreeding ‘rest period’ between mating events (Table 1). Successful spawning events were counted when a female with a recorded mating event during the spawning season deposited an egg mass. Time required for oogenesis between spawning events were parameterized based on observations by Weil & Laughlin [11]. Because females can store viable sperm from a single copulation for several weeks [11, 43], no additional timing requirements were imposed for spawning beyond one prior mating event during the spawning season.

A successful mating event was counted when a male encountered a receptive female. An encounter was defined as the daily paths of two individuals being within the randomly selected perception distance for those individuals. Conspecific perception distance is unknown for queen conch; thus, an informal expert elicitation process was used to parameterize this variable (Table 1). Simulated variability was intended to capture differences in visibility, benthic habitat, and water currents that might carry scent-based cues.

Sexual facilitation was modeled as a positive feedback loop between direct contact or perception of males through chemical cues [33] and receptivity to mating in females [33, 44-46]. Sexual facilitation was modeled as a stochastic process where the likelihood of a female *i* successfully mating at time *t* increased linearly with the number of prior contacts (*C*) with males, up to threshold τ, where mating would be 100% successful:

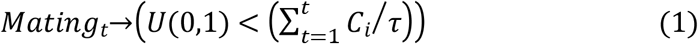

Although sexual facilitation has been demonstrated in other prosobranchs [47-50], it has not been empirically demonstrated in queen conch. Similarly, the accelerated rate of gametogenesis conferred through sexual facilitation is unknown. To encompass these uncertainties, sexual facilitation was run with τ = 0, 5, 10, and 25, respectively, where τ = 0 represents no sexual facilitation.

Percent mating and percent spawning were compared to data from the central Bahamas [30, 32] and the Florida Keys [36]. The data from Delgado & Glazer [36] were filtered to May-July only, to mirror the “peak spawning” season simulated in the model (see Fig 4 in Delgado & Glazer [36]). Goodness of fit from simulation model predictions were compared to logistic regression model fits generated from R package ‘*drc’* using paired t-tests [51]. Simulation model predictions were generated for point observations from field studies through linear interpolation using the *approx* function in R package ‘*stats’* [52].

**Fig 3.**
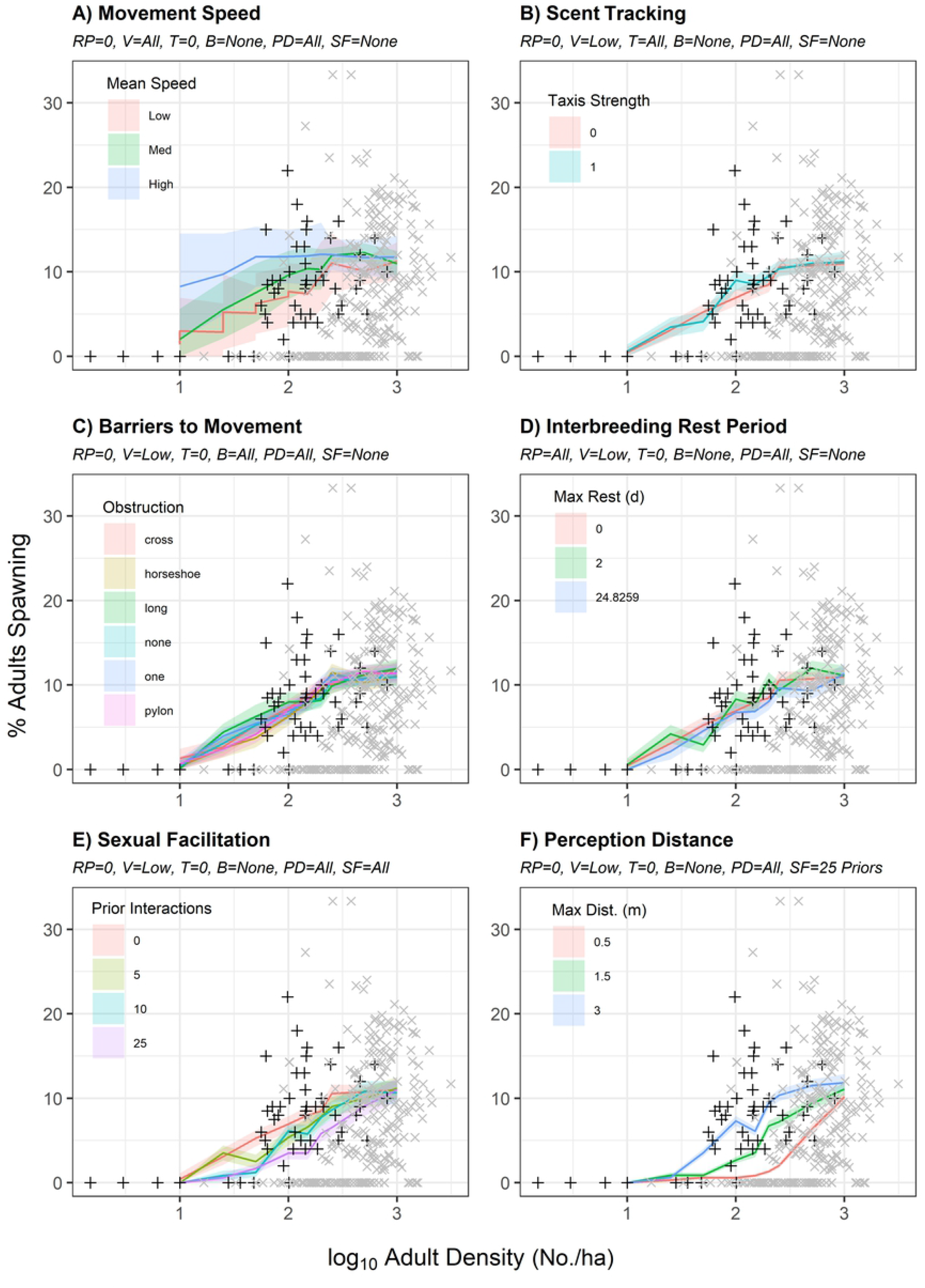
Percent Spawning Relative to Model Parameters. Mean (solid line) and 95% confidence band (shaded ribbon) for percent of simulated adult queen conch successfully spawning relative to log_10_ adult density (N/ha) relative to variation in A) movement speed (V), B) scent tracking (T), C) barriers to movement (B), D) interbreeding rest period (RP), E) sexual facilitation (SF), and F) perception distance (PD). Empirical observations by Stoner & Ray-Culp [30] (black crosses) and Delgado & Glazer [36] (gray x’s) are overlaid for comparison.

**Fig 4.**
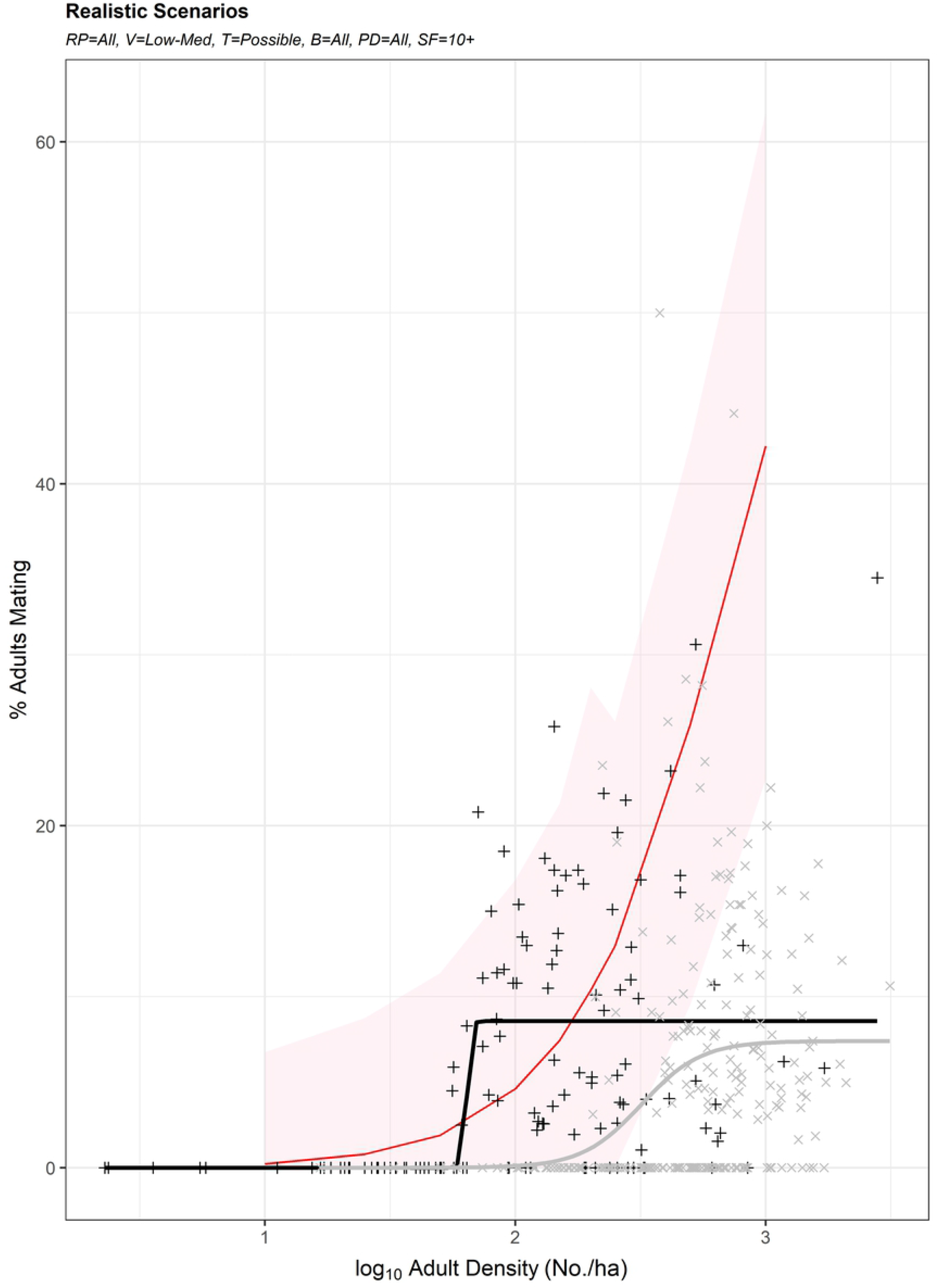
Percent Successful Mating Events Relative to Realistic Scenarios. Mean (solid line) and 95% confidence band (shaded ribbon) for percent of simulated adult queen conch successfully mating relative to log_10_ adult adult density (N/ha) under all interbreeding rest periods, low to medium movement speeds, possible scent tracking, all movement barriers, sexual facilitation with 100% mating success with at least 10 prior interactions, and all conspecific perception distances. Empirical observations by Stoner et al. [32] (black crosses) and Delgado & Glazer [36] (gray x’s) are overlaid for comparison, with corresponding logistic dose-response function fits from R package ‘drc’ [51].

## Results

The influence of density, movement speed, scent tracking, barriers to movement, interbreeding rest period, sexual facilitation, and conspecific perception distance on successful mating and spawning are presented in Figures 2 and 3, respectively. The high movement speed scenario over predicted mating activity at all densities (Fig 2A). All movement speed scenarios provided reasonable fits to empirical observations of spawning activity at densities of 100 or more adults/ha (Fig 3A). Low and medium movement speed scenarios provided better fits to mating activity observations by Stoner el al. [32] as compared to those from Delgado & Glazer [36] (Fig 2A).

At low densities, scent tracking slightly increased mating success; at high densities, it slightly reduced mating success (Fig 2B). Scent tracking had a negligible impact on spawning activity (Fig 3B). Barriers to movement had minimal impacts upon simulated mating (Fig 2C) and spawning activity (Fig 3C). Shorter interbreeding rest periods provided better fits to empirical observations of mating activity by Stoner et al. [32]; longer interbreeding rest periods provided better fits to empirical observations of mating activity by Delgado & Glazer [36] (Fig 2D). Interbreeding rest period did not impact simulated spawning activity (Fig 3D). Sexual facilitation with 100% success at ≥ 10 prior interactions (i.e., τ = 10 or 25) provided better fits to all empirical observations at densities of <100 adults/ha (Figs 2E and 3E). Conspecific perception distance had a large impact upon simulated mating and spawning activity (Figs 2F and 3F); higher perception distances provided better fits to empirical observations by Stoner et al. [32] and Stoner & Ray-Culp [30], lower perception distances provided better fits to empirical observations by Delgado & Glazer [36].

Model simulations for low to medium velocity movement speeds with sexual facilitation requiring at least 10 prior interactions to ensure mating success provided superior fits to mating activity observations by Stoner et al. [32] than logistic regression (Fig 4). In general, simulations over-predicted mating activity relative to observations by Delgado & Glazer [36], and did not account for their repeated observations of mating failure at high densities.

A plot of the relationship between percent of adults mating and percent of adults spawning shows an inflection point at around 25% mating (Fig 5). The modeled relationship based on mechanistic simulations provides a reasonable fit to empirical observations by Stoner & Ray-Culp [30]. The model did not predict the high numbers of spawning observations with no corresponding mating activity observed by Delgado & Glazer [36], but had many outlying observations of higher (>10%) spawning activity at relatively low levels of mating activity (<15%). Simulations suggested a linear increase in total spawning events relative to adult density (Fig 6). Paired t-tests suggested interpolated simulations provided superior fits in 79 (17%) and 2 (<1%) of 457 cases as compared to dose-response curves fit to data from Stoner et al. [32] and Delgado & Glazer [36] data, respectively. All cases of superior fits were realized at low to medium movement velocities. When excluding cases where no mating activity was empirically observed at densities above 100 conch/ha, paired t-tests suggested 104 (23%) and 18 (4%) of 457 mechanistic simulations provided superior fits as compared to dose-response curves fit to the observations of Stoner et al. [32] and Delgado & Glazer [36], respectively.

**Fig 5.**
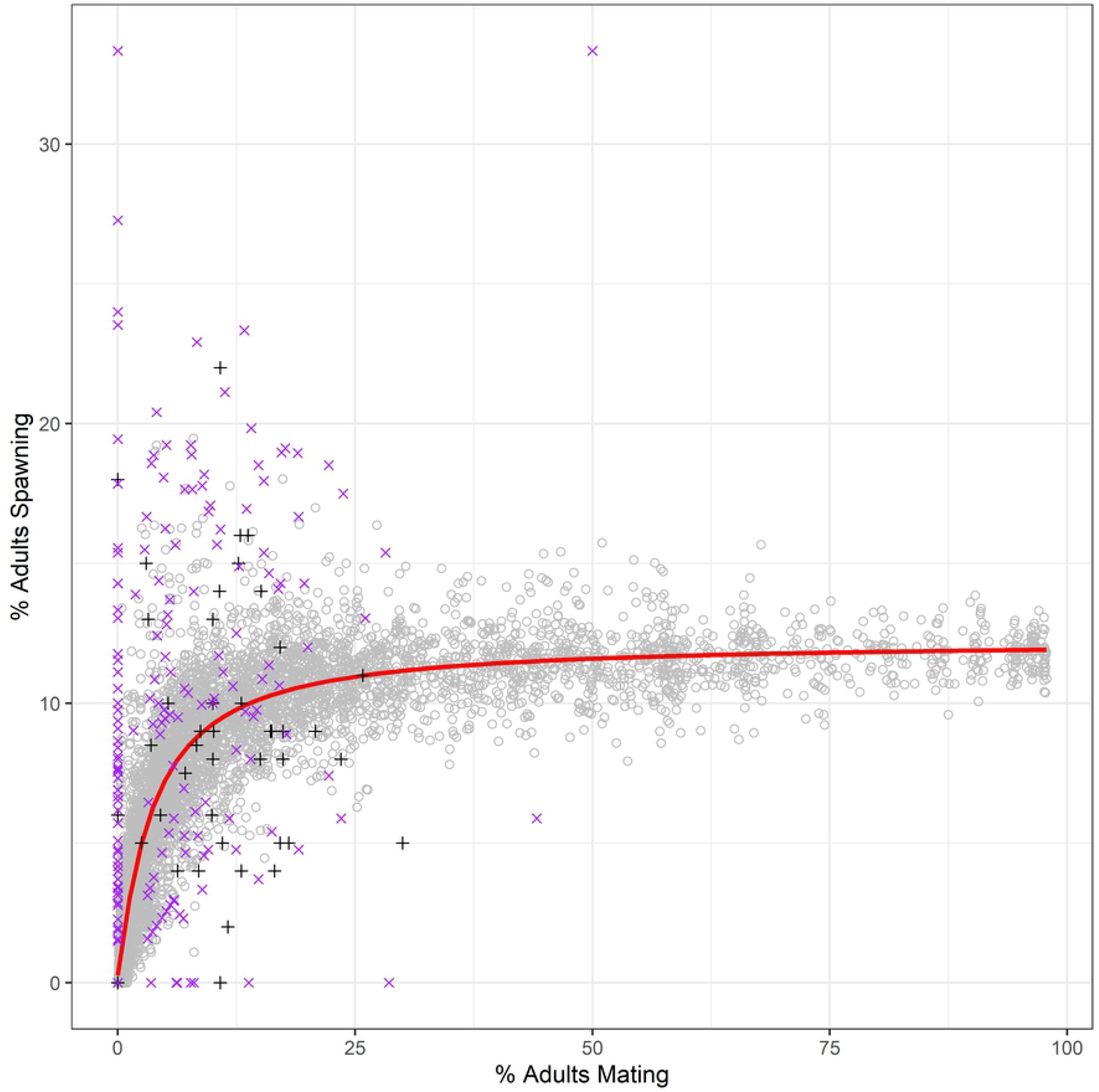
Mating and Spawning Activity. Relationship between mating and spawning activity across all model simulations (gray circles) with logistic dose-response function fit (red line) fit. Empirical observations by Stoner & Ray-Culp [30] (black crosses) and Delgado & Glazer [36] (purple x’s) are overlaid for comparison.

**Fig 6.**
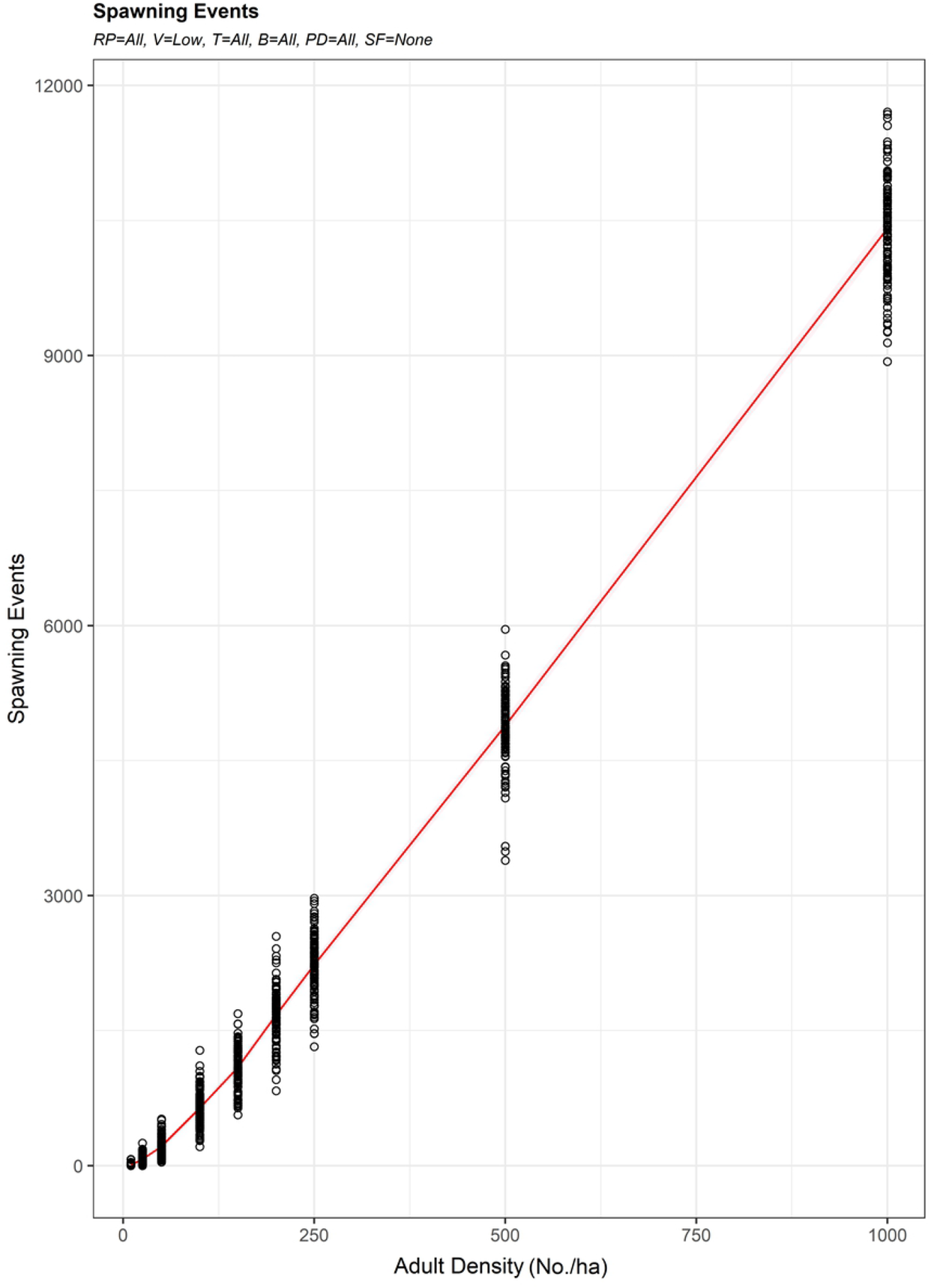
Spawning Events Related to Adult Density. Mean (red line) and all simulated (black circles) spawning events relative to adult density (N/ha).

## Discussion

Depensatory mechanisms have been postulated as a major factor limiting the recovery of overharvested queen conch populations [32, 33]. Reproductive potential is primarily reduced by the removal of spawners from the population [33, 53]. Reproductive potential is secondarily limited by reduced densities, which increase the search time required for encountering potential mates [33]. Our simulations confirm this is especially limiting for slow-moving conch which require internal fertilization for successful mating. This limitation translates directly into limited recovery because the “search time” cost depletes both energy and time resources, meaning gametogenesis will not proceed at maximal rates and thus, populations will not reproduce to their full capacity. Our simulations confirm that limitations on mate finding associated with density is the primary driver behind observed patterns in mating and spawning activity, but similar to field observations by Gascoigne & Lipcius [54], challenges associated with mate finding cannot be the only explanation for lack of reproductive activity at low densities.

Our simulations also indicate that high movement speeds and extensive scent tracking are unlikely explanations for observed trends in queen conch mating and spawning activity (Figs 2A-B and 3A-B). Simulations of these factors provided poor fits to empirical data, and increased levels of movement or taxis would push simulations further from observed trends. Additionally, simulations suggested barriers to movement associated with microhabitat features have little impact on the percentage of the population mating or spawning (Figs 2C and 3C). This is likely because these barriers serve to reflect adult conch back at other conspecifics rather than slowing their rate of movement. This could be a cause for concern with regard to genetic diversity by increasing mating interactions between the same individuals, but does not appear to reduce mating activity. Although interbreeding rest period impacted mating activity in simulations, it did not have a corresponding impact on spawning (Figs 2D and 3D). This is because the longest interbreeding rest period was parameterized according to the interspawning period defined for female conch from Weil & Laughlin [11]. The only satisfactory mechanistic explanation for the absence of empirical observations of mating or spawning activity at low densities was a relatively high requirement (τ ≥ 10) for prior conspecific interactions attributed to sexual facilitation (Figs 2E and 3E). Similarly, the only mechanistic explanation for the reduced mating and spawning activity at all densities observed by Delgado & Glazer [36] relative to Stoner et al. [32] and Stoner & Ray-Culp [30] was reduced conspecific perception distance required for successful mate finding (Figs 2F and 3F).

Queen conch move by anchoring the sickle-shaped operculum against the seafloor and thrusting the foot backward, propelling the shell forward a half body-length at a time [1]. Adults move at varying speeds throughout the year and are capable of extensive seasonal migrations to and from historic spawning grounds. As adults migrate to these spawning areas aggregations are typically formed and individual movements slow when mating and spawning activities begin [5]. Conch may move shorter distances as needed to forage or actively locate compatible mates, but typically remain within smaller areas until returning to their feeding grounds at the conclusion of the reproductive season [5, 16]. Our model simulations using low (2.5 m/d) and medium (4.17 m/d) within-aggregation movement speeds more closely followed field observations of mating activity compared to high (11.36 m/d) movement speeds. The high movement speed used from Doerr and Hill [15] was compiled from an extensive tracking period and included combinations of fine-scale daily movements and large-scale reproductive migrations; however, it did not include estimates of within-aggregation movement rates. Slower movements within aggregations would seem to facilitate higher frequencies of mating activity; at higher densities individuals would not need to travel far to locate a receptive mate. Likewise, easy access to sufficient food supply within the aggregation area would eliminate the need for short foraging trips and allow conch to continue to mate and spawn at high frequencies. Reducing movements to minimize excess energy expenditures during mating and spawning would help to ensure maximum reproductive output, particularly for females, in the form of high overall egg production.

Because queen conch do not move using a slime trail, scent-trail following would presumably be limited in spatiotemporal scope. Our simulations suggested that one-day duration scent-trail tracking out to near the maximum daily movement distance would slightly increase mating activity at lower densities. Ecologically, this could be explained as an increased efficiency in mate finding offsetting the slow movement speed of reproductive adults. Our simulations also suggested that scent-trail following at higher densities might actually lead to a slight reduction in mating activity. This result could be explained by the inability to focus on and track a single individual, leading to inefficiencies in the movement path. This is similar to the well documented “confusion effect” for predatory fish targeting individuals within large schools of fish [55].

Conch can be confined by ecological barriers such as fragmented habitats, the presence of extensive bare sand plains that lack food resources, or areas that may expose them to potential desiccation or anoxic conditions [56, 57]. Natural barriers to movement can serve to isolate populations through suppressed immigration of juvenile and adult conch. For example, in the Florida Keys, the East Harbour Lobster and Conch Reserve in South Caicos, and certain areas of Lee Stocking Island, Bahamas, conch are separated from surrounding habitats by coral ledges, sand bars, and offshore reefs, respectively [5, 56, 58]. In simulations, barriers functioned to limit dispersal, but also increased random interactions between nearby individuals. Habitats with a high degree of reflective barriers could actually serve to increase mating rates in our simple simulations by increasing the probability of repeated encounters with the same individuals. In the wild, this type of environmental bottleneck might cause concerns for genetic diversity within the population, although many factors would come into play that were not evaluated in our simulations [59].

Simulations suggested that longer interbreeding rest periods could play an important limiting role in reproductive success. It is possible that female receptivity to mates will be lower during oogenesis. In high-density aggregations, mating concurrent with egg laying is not uncommon [30]; however, after spawning, females might not attract mates or might avoid mates, creating a rest period. Delays or rest periods in female receptivity are probably not explained by a bioenergetic need to forage after a mating event, as conch have been observed foraging while mating [30, 58]. However, the developmental period between egg masses may have a bioenergetic link, as the development and deposition of large egg masses is an energetically costly event requiring either substantial body reserves or additional energy intake through foraging to be repeated [11, 60]. Overfishing of conch on more productive shallow-water habitats may drive population reproductive potential to deeper, less productive habitats, which may further reduce reproductive output due to increased recovery time between spawning events due to lower habitat quality [8, 61].

In addition to the direct removal of spawners and the increased mate encounter times caused by lower densities, a third potential depensatory mechanism is the breakdown of a positive feedback loop between contact with males and the rate of gametogenesis and spawning in females [33, 44-46]. When reproductive fitness declines such that per capita population growth rate becomes negative, localized extinction may result [62, 63]. This sexual facilitation could be accomplished through direct contact or chemical cues [33]. Copulation in conch is more likely to occur in spawning than non-spawning females, providing an additional positive feedback mechanism that amplifies the effect at high densities [64]. Our model provides mechanistic confirmation that the reductions of densities caused by overharvesting of spawning aggregations increases the probability of recruitment failure beyond what would be anticipated from delays in mate finding alone. This is consistent with field experiment findings from Gascoigne & Lipcius [54], which indicate that in addition to depensatory mechanisms associated with mate finding, delayed functional maturity at low-density sites can explain declines in reproductive activity. As such, understanding depensatory thresholds seems absolutely critical to effective fisheries management [36].

Adult density had the largest effect on mating and spawning activity. Sexual facilitation was the only mechanistic explanation for extremely low (or lack of) mating rates observed at low densities. Similarly, perception distance was a major controlling factor of mating and spawning rates at higher densities. When queen conch were assumed to have very limited (max of 0.5 m) perception distance for mating encounters, simulation outputs more closely patterned empirical observations in the Florida Keys back reef by Delgado & Glazer [36]. When queen conch were assumed to have fairly broad (up to 3 m) perception distance, simulation outputs more closely patterned empirical observations in Bahamian waters by Stoner et al. [32] and Stoner & Ray-Culp [30]. Ecologically, perception distance could be interpreted as near-field ability to visually or chemically locate potential mates. In the field, perception distance might vary based on the strength and duration of chemical cues in the water or on the substrate, the direction and strength of current flows between potential mates, and water clarity. Inferring from studies with other gastropods, queen conch likely detect conspecifics and predators through their chemosensitive tentacles and use their keen eyesight to orient subsequent movements [65]. The eyes of *Strombus/Lobatus* are among the best developed of those found in gastropods [66], and it is likely that conch can converge on objects during visual fixation [67]. We are unaware of any studies of how far queen conch can see, but our simulations suggest some of the differences in mating activity observed between Stoner et al. [32] and Delgado & Glazer [36] could be attributed to differences in perception distance. It is possible that the clear waters and relatively flat, shallow habitats of the Bahamas provide a greater perception distance (closer to 3 m) than the rugose, lower-visibility back-reef sites surveyed by Delgado & Glazer [36]. Further studies on the conspecific perception distance and visual acuity of queen conch are needed to validate this hypothesis.

Our model did not account for phenotypic differences between mature adult conch. Although the long-standing assumption has been that variation in shell morphology results from genetic differentiation, for queen conch phenotypic differences are more likely induced by localized ecological conditions. Growth is negatively related to depth [68] due to reduced light availability, indirectly affecting both the abundance and quality of food available [5]. Samba conch are a phenotypically smaller variant of queen conch and have been reported throughout the Caribbean [4, 69, 70]. They are thought to have lower fecundity due to their smaller size, which limits space for gonadal tissue [71]. Evidence supporting limited food resources as a major driver in phenotypic differences has been reported from high-density populations in marine reserves, where individuals grew slower and exhibited thickened shell material, likely as a result of intra-specific food competition [4, 56]. It is currently unknown whether these somatic varieties experience reduced fecundity, and therefore reproductive output; however, the possibility exists that increased shell thickness in females limits the internal space available for development of the ovaries.

Our simulations assumed an environmentally consistent 92-d peak spawning period. Environmental stochasticity might explain some of the variance observed in the field. In marine species, including mollusks, environmental triggers including rapid changes in water temperature or the detection of conspecific gametes in the water are often implicated in the initiation of gametogenesis and reproductive activities [73]. For queen conch, multiple studies have identified increasing water temperature and photoperiod as the stimulus for reproductive migrations and the subsequent initiation of mating [5, 11], and recent evidence has verified the presence of sex hormones in conch feces [74]. Concentrations of estrogen, progesterone, and testosterone increased in conjunction with each phase of the conch reproductive season, indicating that these hormones are linked to the reproductive process [74]. Active hormone detection by conspecifics would positively influence encounter rates of low-density populations and could explain our model outcomes for scent-tracking simulations where mating success increased at lower densities.

Increasing water temperature due to climate change is likely to alter the timing and duration of the queen conch reproductive season. In warmer regions, conch have been observed mating and spawning year round [1, 75]; however, reproduction can also cease as temperatures approach 31°C [76]. Increasing water temperatures may initially extend the reproductive season and shift peak mating and spawning periods, but further increases may subsequently shorten the season as temperatures reach a threshold. If adult conch respond to temperature increases by moving from shallow mating grounds to deeper waters with potentially diminished habitat quality, overall reproductive output may decrease.

In addition to environmental drivers that influence reproductive success, there are biological factors that can negatively impact individual reproductive output. Histological examination of digestive gland samples collected from queen conch throughout the majority of their Caribbean range revealed the presence of a coccidian Apicomplexan inclusion body [77]. A higher abundance of these inclusion bodies appeared to correspond with reproductive abnormalities such as reduced frequency of gametogenesis, delayed maturity, and low gonad activity and spawn stages [78, 79]. However, recent studies have suggested that these inclusion bodies are not parasitic and the individuals sampled appeared to be reproductively healthy [80, 81]. This recent finding warrants further research into overall reproductive impacts, particularly for adult conch at reduced density levels. This could be useful in refining our model to further examine reproductive impacts in populations at seemingly adequate adult densities but exhibiting reduced overall spawning activity.

This model may be useful in estimating the numbers of eggs produced by spawning aggregations at given densities. The predicted number of spawning events increased exponentially with adult density. Delgado & Glazer [36] found egg masses contained a mean of 364,532 (SE = 33,383) eggs. Our model estimates over 200 million eggs would be produced within a single hectare over the course of a spawning season with 100 adults, with an increase of over an order of magnitude anticipated with densities approaching 1000 adults/ha. Our model was intended to simulate reproductive dynamics during a 92-d peak spawning season. As such, the number of eggs laid would be an underestimate relative to many areas where the spawning season is more protracted. Simulating only the peak spawning season avoided the need to parameterize increasing concentrations of sex hormones and associated rates of oogenesis with the progression of the spawning season [42] or changes in density and movement rates associated with the formation and dissolution of the aggregation.

This model may also be useful for identifying reference points that avoid recruitment failure. Cross-shelf density thresholds for mating and spawning reported by Stoner & Ray-Culp 30 in the Bahamas were 56 adults/ha for mating and 48 adults/ha for spawning. Similarly, Stoner et al. [32] report threshold densities for mating of 47 to 74 adults/ha. By contrast, Delgado & Glazer [36] report aggregation density thresholds of 204 adults/ha for mating and 90 adults/ha for spawning, respectively. These discrepancies may be partially explained by differences in methodology. Stoner & Ray-Culp [30] recorded reproductive behaviors for conch outside the survey circles in which they estimated density, and individuals were counted as mating if they were in mating position but not actually copulating [36]. The United Nations Environment Programme (UNEP) has recommended a reference point of 100 adults/ha to avoid impacts to recruitment [82]. Recent studies [36] and practical application in Jamaica [83] have suggested this threshold may be insufficient to avoid population collapse. Our simulations suggested an inflection point with approximately 25% of the population mating resulting in near-peak reproductive potential, as measured by percent spawning (Fig 5). Simulations incorporating sexual facilitation (Figs 2E-F and 3E-F) suggest densities >200 adults/ha are necessary to achieve high levels of spawning output. Furthermore, simulations suggest that if sexual facilitation is required and perception distance is minimal, meaningful spawning will not occur at densities <250 adults/ha (Fig 3F).

For queen conch and similar motile invertebrates that must locate conspecifics for reproduction, population density is one of the most critical factors in maintaining the reproductive output of the stock. However, other drivers of mating and spawning success examined in our model simulations indicate that population density is not the only factor to consider. Our simulation results suggest that biological characteristics of queen conch such as scent-tracking ability, rest period, sexual facilitation, perception distance, and movement speeds interact with density to varying degrees to influence mating and spawning frequencies, and thus, total reproductive output. Further modelling exercises incorporating refined biological parameters or additional environmental drivers can potentially be used to guide the development of innovative management strategies and enhance conservation efforts.

## Acknowledgments

Thanks to Allan Stoner, Robert Glazer, Gabriel Delgado, Calusa Horn, and Mandy Karnauskas for valuable insight during the development of these simulations.

## Supporting information

**S1 Data. Complete set of model parameters and input values used in model simulations**. (XLSX)

**S2 File. R code for the queen conch reproduction simulations**. (R)

